# The effect of environmental enrichment on social dominance and welfare in a cichlid fish

**DOI:** 10.64898/2025.12.11.693693

**Authors:** Farjana A. Chamily, Benjamin R. Vanderklok, Olivia D. K. Buzinski, Calvin Ketterl, Zachary D. Hager, Ronald G. Oldfield, Jonathan D. Kelty, Peter D. Dijkstra

**Author notes:** <u>Corresponding author:</u> Peter Dijkstra.

## Abstract

Environmental enrichment can have complex, contradictory effects on aggression and animal welfare. Although increasing enrichment may reduce aggression by limiting the visibility of competitors, it may also intensify territorial defense and harm the welfare of subordinates who are the target of territorial aggression. However, it remains unclear how variation in the complexity of an enrichment structure defended by a single dominant individual influences welfare outcomes for subordinates. In the cichlid fish *Astatotilapia burtoni*, dominant males defend structures as mating territories, whereas subordinate males do not. To establish social hierarchies, we housed two differently sized males (larger become dominant) and six females in one compartment containing one defendable structure. We manipulated structure complexity by placing 1, 2, or 3 halves of terracotta pots clumped together. Increasing cave number did not enhance territoriality: dominant males showed similar aggression rates, relative gonad sizes, and testosterone levels across treatments. Cave number did not significantly affect mortality, body condition, growth rate, or fin damage in subordinate males. Although relative gonad size and testosterone levels were generally higher in dominants, these differences were not always significant in the 2 and/or 3-cave treatments, suggesting weaker physiological differentiation between social states. The behavioral data supported this pattern, with status-specific differences in chase rates declining with increasing cave number. Overall, increasing environmental enrichment had little effect on welfare, but resulted in reduced physiological and behavioral distinctions between dominant and subordinate males. Therefore, careful consideration of enrichment strategies is essential for accurately interpreting status-specific outcomes in laboratory settings.

## Introduction

Many animal species form social hierarchies and aggressively defend their rank or limited resources by showing hostile behaviors towards cohabitants. These behaviors can lead to a decreased quality of life for captive organisms and may skew results in research settings (du Sert et al., 2020; Theil et al., 2020). In a natural environment, there is typically a greater opportunity for individuals at the receiving end of aggression to escape and honor territory boundaries set by individuals with higher social rank. In a laboratory cage or a zoo enclosure, opportunities to avoid conflict are greatly reduced, potentially resulting in lower welfare in subordinate individuals that are the target of aggression. For example, mice that are kept in a laboratory setting cannot escape intraspecific aggression from dominant individuals, and consequently, intra-cage aggression is potentially the most common source of morbidity in mouse facilities (Lidster et al., 2019; Theil et al., 2020).

Environmental enrichment is a common tool to improve welfare by allowing animals to express their natural behavioral repertoire. It may also mitigate the negative consequences of aggression by providing opportunities for hiding to avoid conflict (Benefiel et al., 2005; Kaliste et al., 2006; Young, 2013; Stevens et al., 2021). For example, partial cage dividers, used to increase environmental complexity, reduced aggressive events in male mice, bite wounds, anxiety, and improved overall welfare by creating more opportunities to avoid conflict (Tallent et al., 2024). Physical complexity may create barriers between competitors, which can lower intraspecific aggression levels. For instance, in brown trout (*Salmo trutta*), complex habitats using woody debris decreased opportunities for aggressive interactions by visually isolating fish, promoting animal welfare as indicated by reduced mass loss (Sundbaum and Näslund, 1998). Structural complexity may help delimit individual territories within a captive setting, allowing subordinate individuals to respect territorial boundaries and avoid conflict (Galhardo et al., 2008; Smith, 2011). However, physical structures used to enrich the environment can raise resource value and encourage active defense by the owner against competitors, as shown in mice, zebrafish, and cichlids (Kaliste et al., 2006; Dijkstra et al., 2008; Mandrekar and Oldfield, 2008; Wilkes et al., 2012; Giles et al., 2018). Although territorial aggression can be used as a way to induce physiological stress in subordinate individuals (Patki et al., 2013; Ibi et al., 2017; Lehmann et al., 2019; Snyder-Mackler et al., 2020), strategies to reduce the amount of aggression are vital for both animal welfare (through reducing pain and suffering) and high-quality scientific research.

The conflicting effects of enrichment on aggression and animal welfare may be due, in part, to increased environmental heterogeneity supporting a higher density of territorial individuals (Blanchet et al., 2006; Eason and Stamps, 1992) or more equal distribution of limited resources (Basquill and Grant, 1998). For example, in rainbow trout (*Oncorhynchus mykiss*), territorial enrichment using cobbles and dividers resulted in smaller individual territories as habitat visibility decreased (Imre et al., 2002). These changes in the density of territorial individuals or the degree of resource monopolization may greatly impact welfare by altering the amount of escalated aggression directed at subordinate individuals (Ruberto et al., 2024). Consequently, enrichment-induced changes in the dominance hierarchy make it difficult to isolate the mechanisms that modulate the amount of territorial aggression and opportunities for conflict avoidance. Hence, clarifying the effect of environment enrichment on animal welfare should be carried out in controlled settings where the density of dominant individuals is kept constant across enrichment treatments. Specifically, it is currently unclear how variations in the complexity of a compact enrichment structure defended by a single dominant individual influence the welfare outcomes of subordinate individuals.

The African cichlid fish *Astatotilapia burtoni* exhibits a social hierarchy in which dominant males are brightly colored, aggressively defend territories used to attract females for spawning, and have an upregulated reproductive system as indicated by large gonads (Maruska and Fernald, 2018). Subordinate males are not brightly colored, do not defend a territory, and are reproductively inactive. In many *A. burtoni* studies, dominant and subordinate males are studied relative to their behavior (Piefke et al., 2021), neural regulation of reproduction and competition (Maruska and Fernald, 2011; O’Connell and Hofmann, 2012; Alward et al., 2019), or regulation of oxidative stress (Border et al., 2019; Dijkstra et al., 2024). Previous studies suggest that excessive aggression from dominant males can lead to injury and death in subordinate fish (Alcazar et al., 2016; Alward et al., 2020), but studies on enrichment and animal welfare are lacking in this species.

We tested how environmental enrichment influences markers of social status and welfare in *A. burtoni* mixed-sex groups (2 males, 6 females) using a common housing paradigm in which a single, larger dominant male defends a territory (Fialkowski et al., 2022). To manipulate enrichment, groups were housed with structures consisting of either 1, 2, or 3 halves of terracotta pots (referred to as ‘caves’). In all treatments, each structure could be defended by one dominant male (the two and three cave structures were clumped together). Hence, we were able to assess how animal welfare is impacted by environmental complexity with experimental groups, with each containing one dominant male and several subordinate fish. Since a more complex territory raises resource value, we predicted that increased cave complexity would increase aggressive behavior, testosterone levels, and relative gonad size in dominant males only. Such a hypothesized increase in territoriality in more complex treatments could lead to reduced welfare of subordinate fish because of increased aggression from the dominant male. However, because increasing the number of caves would reduce visibility, create more hiding opportunities, and potentially distract territorial males more (by encouraging more cave inspections), we predicted that increasing cave number would nevertheless improve welfare in subordinate fish as indicated by increased body condition, growth, and fewer fin splits.

## Material and Methods

### Animals

Males and females of *A. burtoni* (aged approximately 8-12 months) were taken from a laboratory-bred population originating from Lake Tanganyika, Africa (Fernald and Hirata, 1979). Fish were raised in 400-L mixed-sex tanks with gravel substrate maintained at 28℃ under a 12-hour light/dark cycle, with gradual light changes to simulate natural dawn and dusk. Fish were fed cichlid flakes (Omega One cichlid flakes) and pellets (Omega One cichlid pellets, 2 mm) between 8:00 and 10:00 am daily. Fish were individually tagged just below the dorsal fin with colored beads attached to a plastic tag using a stainless-steel tagging gun (Avery-Dennison, Pasadena, CA). The study was conducted between 12^th^ of July and 14^th^ of August of 2024 at Central Michigan University in Mount Pleasant, Michigan, USA. All procedures were approved by the Central Michigan University Institutional Animal Care and Use Committee (2021-460).

### Experimental design

At the start of the experiment, both males and females were collected from the same 400-L grow-out tank and measured for standard length and weight. For the experimental setup, several 110-L tanks were each divided with a clear, perforated screen into two 55-L sections to create two compartments (Fialkowski et al., 2021). To assign social status based on size difference, one larger male and one smaller male were placed in each compartment along with 6 females. In this setup, fish were able to interact physically with members of their own group and visually with the others in the adjacent compartment. This allowed for the full expression of social behaviors including aggressive interactions between dominant males between compartments. Halved terracotta pots (14 × 7 × 13 cm) were placed in the corner of each compartment (∼30 cm from the divider; **Figure 1**) to serve as defendable territories. The structural complexity was manipulated by placing 1, 2, or 3 halves of terracotta pots (caves’), with an approximately 2 cm gap between the caves and the tank wall to allow the fish to enter the caves (see Supplementary Figure with detailed cave arrangement). We set up a total of 12 groups in the 1-cave treatment, 13 groups in the 2-cave treatment, and 13 groups in the 3-cave treatment (each containing 2 males and 6 females). Of the 38 groups, five were excluded from the analyses: one due to misidentification of a female as a male (3-caves), one because a female became dominant over the males (3-caves), one where two dominant males emerged (3-caves), and two due to male mortality in the first week of the experiment (3 and 2-caves). A total of 304 fish were used in this study (initial weight and standard length of males [mean ± SE]: 8.16 g ± 1.03 [SE], 58.78 mm ± 1.28 [SE]; females: 3.2 g ± 0.07 [SE], 48.4 mm ± 0.37 [SE]). Only male data were analyzed in this study; female data will be reported elsewhere.

**Figure 1:**
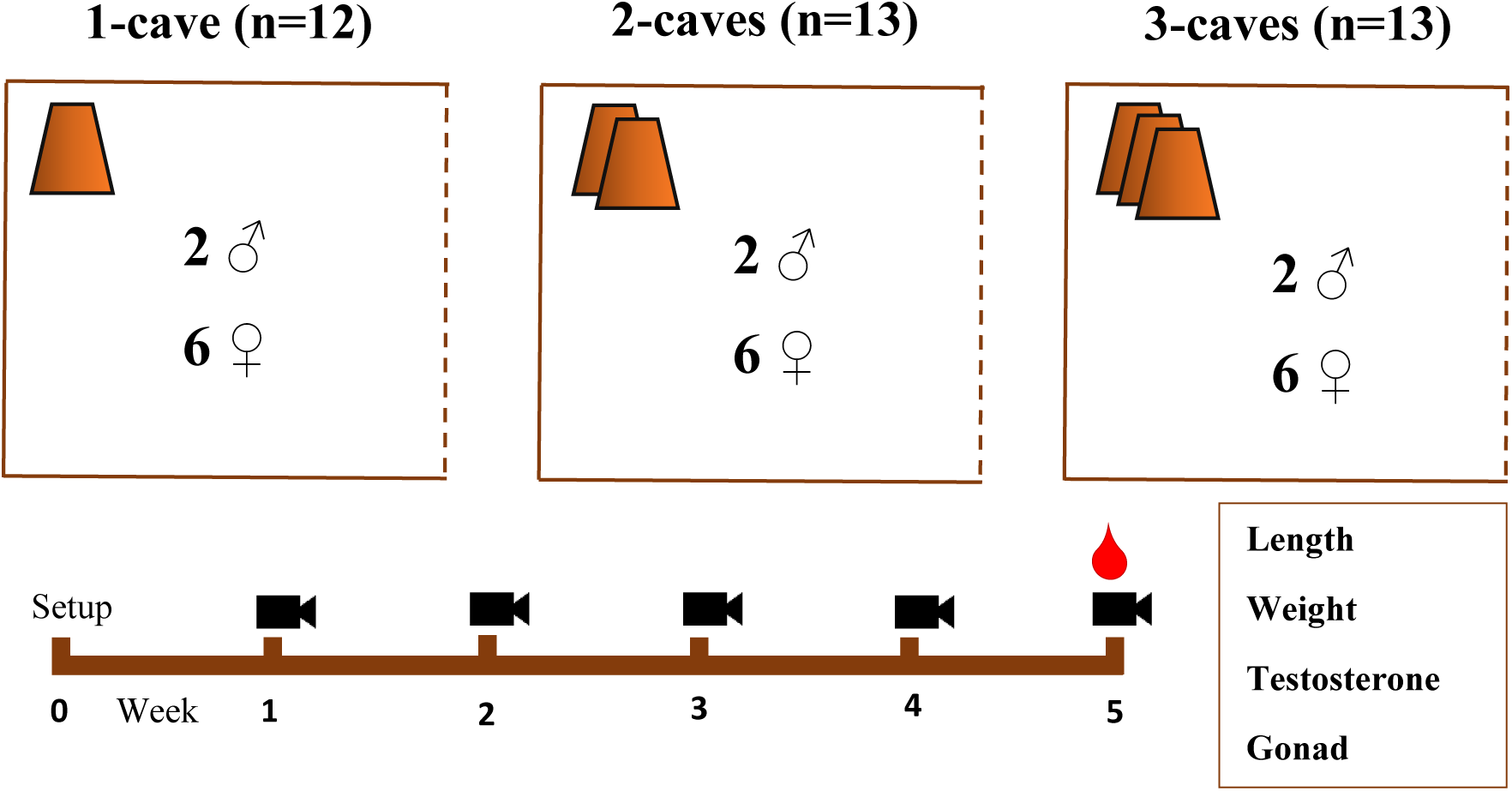
Experimental design. Two males and six females were housed in compartments with a neighboring compartment (also with two males and six females) visible behind a perforated screen (dashed line). The larger male was dominant while the smaller male was subordinate. Halved terracotta pots (‘caves’) were placed in the corner of the tank in the 1-cave (n = 12), 2-caves (n = 13) and 3-caves (n = 13) treatments. Tanks were filmed for 5 minutes and 30 seconds once per week. After the final filming session, males were measured and sacrificed for tissue collection. Blood and gonads were collected from each male to assess physiological markers of social status.

### Behavioral and morphological analysis

After introducing the fish into each compartment, the dominance hierarchy was typically established in one to two days, with the larger male gaining social dominance and the smaller male being forced into the subordinate position in all groups that were included in the analysis (n = 35). After a 1-week acclimation period, we recorded social status and injury by quantifying fin splits for all fish 3 times per week.

Dominant males displayed bright coloration, defended territories, and exhibited a distinct dark eye bar, whereas subordinate males had cryptic coloration and shoaled with females. As an indirect measure of aggression, we quantified fin damage, as the number of splits in the dorsal, pectoral, anal, and caudal fins (modified from Hoyle et al., 2007; Noble et al., 2008; Oldfield, 2011).

For the next 5 weeks, once per week before 10 a.m. fish were filmed (Canon EOS 70D) for 5 minutes and 30 seconds for behavioral quantification, with the final recording made on the morning of tissue collection (Fialkowski et al., 2022). Using the videos, an observer scored the frequency of behaviors as previously described (Border et al., 2019; Fialkowski et al., 2021). As social dominance behaviors, we quantified the number of chases (defined as a rapid movement of one fish towards another, with the target typically fleeing), display behaviors (such as border threat and lateral display), cave visits, and courtship displays. We also scored the frequency of fleeing behaviors and foraging behaviors.

### Tissue collection

At the end of week five, following the final behavioral observation, both males and females were measured for standard length (SL; mm) and weight (g). Blood (25-100 μL) was collected from all males. Male gonads were extracted and weighed to calculate the gonadosomatic index (GSI) (Fialkowski et al., 2022).

Blood was drawn from the caudal vein using heparinized 26-gauge butterfly needles (Terumo) and placed in heparinized microcentrifuge tubes, which were kept on ice until centrifugation at 4000 g for 10 minutes to separate the plasma. The plasma was then aliquoted for assays and stored at −80°C.

### Circulating Testosterone

We measured testosterone levels from 7 µL plasma samples using a competitive-binding ELISA kit and protocol (Enzo Life Sciences), as previously described (Border et al., 2019). We measured optical density at 405 nm using a Tecan M200 Infinite plate reader. We reported the values as nanograms of testosterone per milliliter of plasma (ng/mL). The intra-assay coefficient of variation (CV) was 1.3% and the inter-assay CV was 10.7%.

### Statistical analysis

All analyses were performed in R v4.30. Relative gonad mass (gonadosomatic index [GSI = (gonad mass/body mass)*100]) was used as an indicator of upregulation of the reproductive system and territoriality (Bolger and Connolly, 1989). Body condition was calculated using the formula [((body mass/ (standard length)^^3^) * 100)] (Bagenal and Ricker, 1978) and specific growth rate was calculated using [((log(final weight)-log(initial weight))*100)/34] (Jobling, 1983). To assess differences in GSI, testosterone, specific growth rate, and body condition based on social status, we fitted linear mixed models (LMMs) that included social status (dominant or subordinate) as a fixed factor and group ID as a random factor, using the lmer function from the “lme4” package (Bates et al., 2015; Culbert et al., 2023). We then examined whether these markers were associated with cave treatments by fitting LMMs that included cave number, social status (dominant or subordinate), and their interaction as fixed factors, with group ID as a random factor. To investigate how social behaviors (such as chases, aggressive displays, courtship, cave visits, and fleeing behaviors) were affected by cave treatments, we employed generalized linear mixed models (GLMM) with status as the independent variable. GLMMs were fitted using negative binomial distribution using the glmmTMB package (Brooks et al., 2017). We analyzed dominant and subordinate males separately for all analyses assessing the relationship between social behaviors and hormone levels, as the distributions of these variables were highly divergent between the groups. We used two-tailed tests. We report mean ± SE of all model estimates.

## Results

### 1. Effects of enrichment on animal welfare Animal injury and mortality

We observed the mortality of two subordinate males at the onset of the experiment before filming began; however, no further male mortality was found throughout the study period. One of the males that died was from the 2-caves treatment, and the other was from the 3-caves treatment. In terms of injury, subordinate males had more fin splits compared to dominant males (GLMM, status: −1.726±0.112, z= −15.333, *p* < 0.001) in all cave treatments. Within subordinate males, the number of fin splits increased with time (GLMM, week: 0.115±0.043, z= 2.664, *p* = 0.008), but effect was not influenced by treatments (GLMM, caves * week: −0.068±0.053, z= −1.297, *p* = 0.194) (**Figure 2c**). When fitted as main effect, treatment did not affect the number of fin splits (GLMM, caves: −0.089±0.1, z= −0.884, *p* = 0.377) after controlling for the effect of status (GLMM, status: −1.726±0.112, z= −15.333, *p* < 0.001).

**Figure 2:**
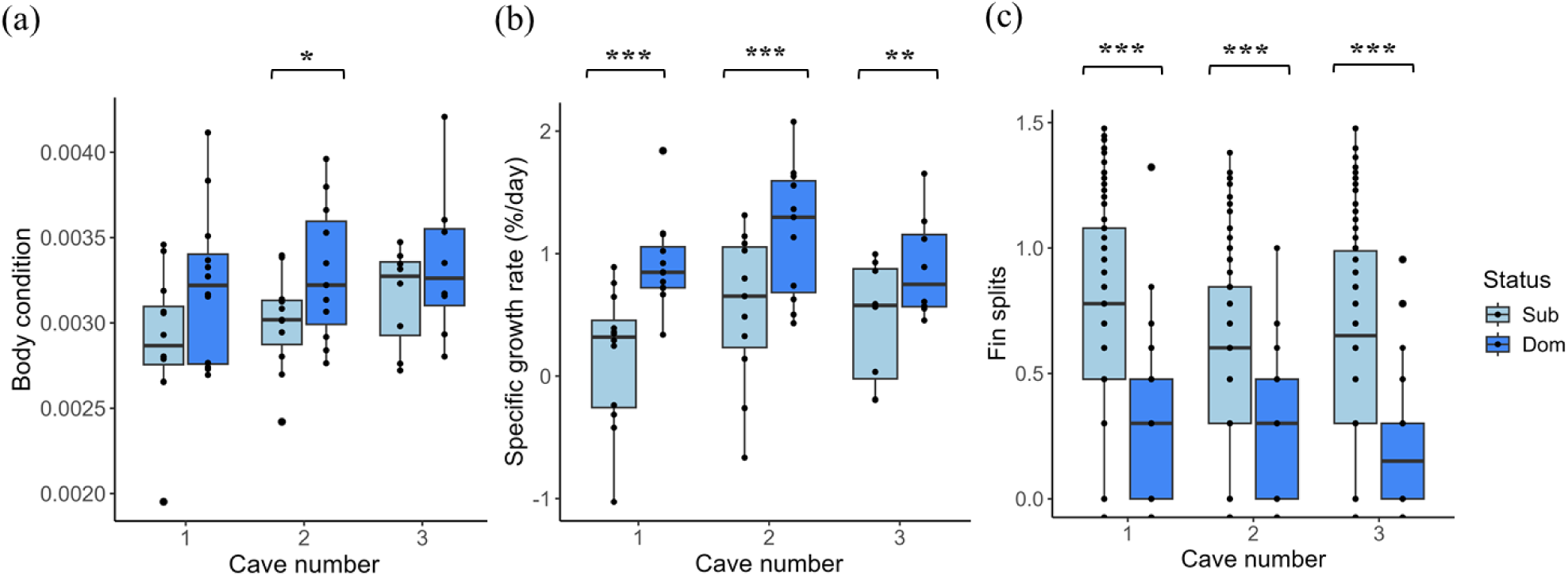
(a) Body condition, (b) specific growth rate, (c) fin splits (log-transformed) as markers of welfare of subordinate (Sub) and dominant (Dom) males for 1-cave, 2-caves and 3-caves treatments. Boxes represent 25th to 75th percentiles. Error bars represent data range. \**p* < 0.5, \*\**p* < 0.01, \*\*\**p* < 0.001.

#### Body condition

In subordinate males, cave number had a positive effect on body condition (LM, caves: 0.0001±0.00007, t = 1.675, **Table 1**), but this effect was not significant (*p* = 0.106, **Table 1**). In dominant males, body condition did not vary significantly with cave numbers (LM, caves: 0.00006±0.00009, t = 0.643, *p* = 0.526). Within each treatment, dominant males had a significantly higher body condition than subordinate males in the 2-caves treatment (LMM, status: −0.0003±0.00009, t_11.8_ = −3.04, *p* = 0.0105), but this status difference was not significant in the 1-cave treatment (LMM, status: −0.0003±0.0001, t_13.1_ = −1.863, *p* = 0.085) or the 3-caves treatment (LMM, status: −0.0002±0.0001, t_9.14_= −1.343, *p* = 0.212) (**Figure 2a**). As expected, dominant males exhibited higher body condition than subordinate males (*p* = 0.004, **Table 1**). However, this effect did not vary with cave number treatments (*p* = 0.574, Table 1). After controlling the effect of social status, cave number treatments were found not to have an effect on body condition (*p* = 0.137, **Table 1**).

**Table 1:** Relationships between markers of welfare (body condition, specific growth rate) and social status (gonadosomatic index and testosterone) with cave treatments in dominant and subordinate *Astatotilapia burtoni* males. We analyzed the data using linear mixed models with social status as a fixed factor and group identity as a random factor. Shown are the interaction effects and main effects. Significant effects (*p* < 0.05) are indicated in bold.

| Marker | Effect | Estimate | df | t value | p value |
| --- | --- | --- | --- | --- | --- |
| Body condition | Cave | 0.00009±0.00006 | 30.11 | 1.526 | 0.137 |
|  | Status | 0.0003±0.00009 | 30.92 | 3.103 | **0.004 \*\*** |
|  | Cave * Status | -0.00006±0.0001 | 29.93 | -0.569 | 0.574 |
| Specific growth rate (%/day) | Cave | 0.085±0.102 | 31.37 | 0.832 | 0.412 |
|  | Status | 0.625±0.082 | 30.80 | 7.625 | **<0.001\*\*\*** |
|  | Cave * Status | -0.157±0.098 | 30.14 | -1.597 | 0.121 |
| Gonadosomatic index | Cave | 0.033±0.030 | 30.18 | 1.085 | 0.287 |
|  | Status | 0.390±0.040 | 30.34 | 9.68 | **<0.001\*\*\*** |
|  | Cave * Status | -0.019±0.051 | 29.47 | -0.387 | 0.701 |
| Testosterone (ng/mL) | Cave | 2.576±1.941 | 55 | 1.327 | 0.19 |
|  | Status | 14.895±3.091 | 55 | 4.819 | **<0.001\*\*\*** |
|  | Cave * Status | -5.266±3.821 | 55 | -1.378 | 0.173 |

#### Specific growth rate

Specific growth rate was higher in dominant males compared to subordinate males (*p* < 0.001, **Table 1**) as expected. However, this effect did not vary across cave number treatments (*p* = 0.121, **Table 1**). When comparing each cave treatment, dominant males exhibited significantly higher SGR than subordinate males in the 1-cave treatment (LMM, status: 0.757±0.167, t_13.1_ = −4.531, *p* < 0.001), 2-cave (LMM: status, −0.607±0.109, t_11.4_ = −5.558, *p* < 0.001), and the 3-caves treatments (LMM, status: −0.441±0.125, t_9.14_ = −3.528, *p* = 0.006) (**Figure 2b**). After controlling the effect of social status, SGR of males did not vary with cave treatments (*p* = 0.412, **Table 1**). Cave treatment did not affect SGR for dominant males (LM, caves: 0.006±0.105, t = 0.056, *p* = 0.956) or for subordinate males (LM, caves: 0.161±0.129, t = 1.249, *p* = 0.222).

### 2. Effects of habitat complexity on markers of social status Gonadosomatic index

As expected, gonadosomatic index (GSI) was higher in dominant males than in subordinate males (*p* < 0.001, **Table 1**). This effect did not vary significantly across cave number treatments (*p* = 0.701, **Table 1**). Within each cave treatment, dominant males exhibited significantly higher GSI in the 1-cave treatment only (LMM, status: −0.442±0.078, t_13.1_ = −5.612, *p* = 0.001), but not in the 2-caves treatment (LMM: status, −0.164±0.157, t_12.6_ = −1.048, *p* = 0.314) or the 3-caves treatment (LMM, status: −0.788±0.352, t_9.14_ = −2.241, *p* = 0.0513) (**Figure 3a**). Cave number treatment did not have a significant effect on GSI (*p* = 0.287, **Table 1**) after controlling for the effect of social status (*p* < 0.001, **Table 1**). Within social states, cave number treatment did not significantly influence GSI for dominant (LM, caves: 0.193±0.125, t = 1.547, *p* = 0.133) or subordinates males (LM, caves: 0.053±0.075, t = 705, *p* = 0.486).

**Figure 3:**
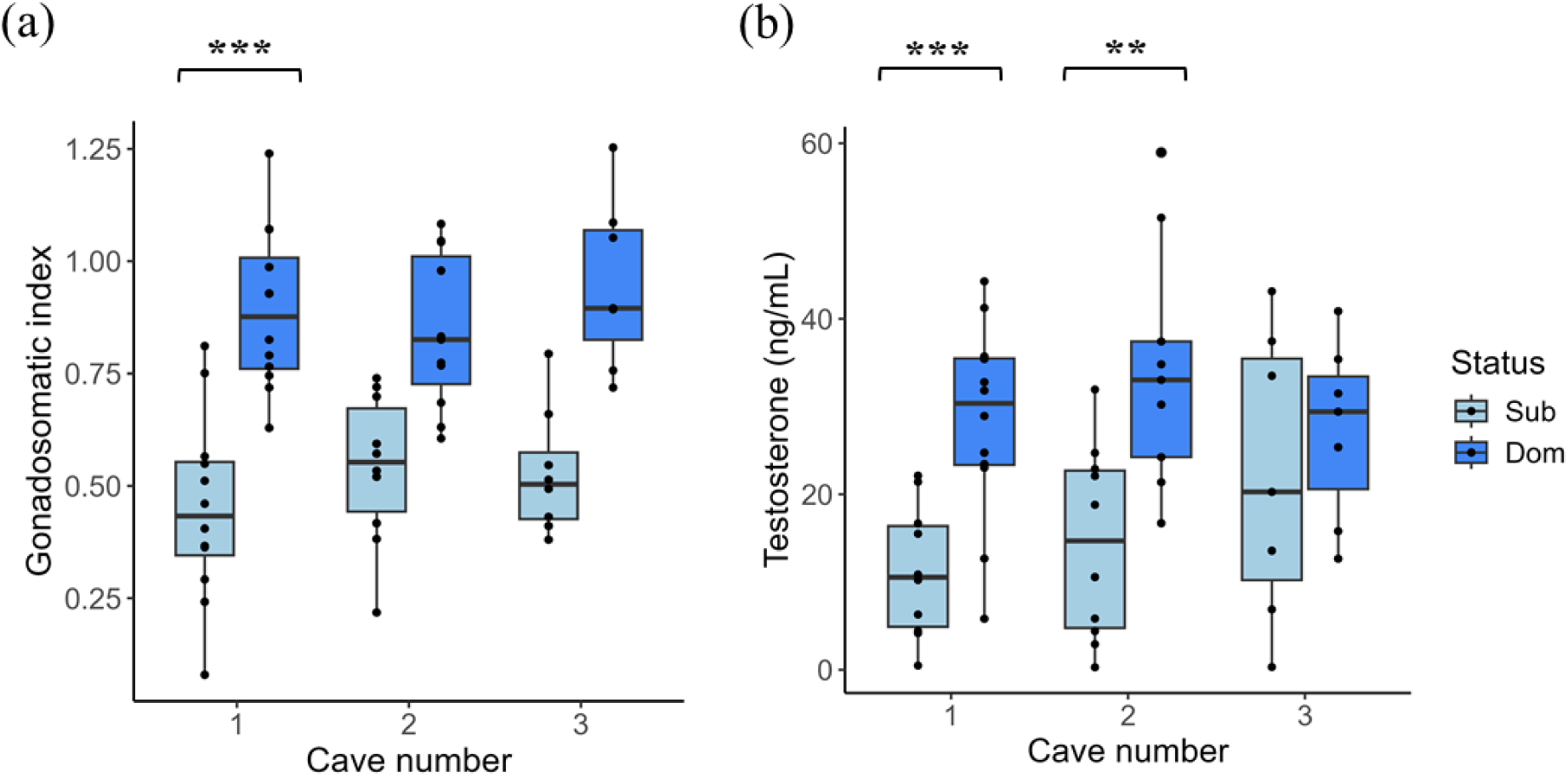
(a) Gonadosomatic index and (b) testosterone levels as markers of social status of subordinate (Sub) and dominant (Dom) males for 1-cave, 2-caves and 3-caves treatments. Boxes represent 25th to 75th percentiles. Error bars represent data range. \*\**p* < 0.01, \*\*\**p* < 0.001.

#### Testosterone

Dominant males exhibited significantly higher testosterone levels compared to subordinates (*p* < 0.001, **Table 1**). This effect did not vary significantly with cave number treatments (LMM, social status * caves: *p* = 0.173, **Table 1**). Dominant males showed significantly higher testosterone levels in the 1-cave (LMM, status: −17.1±4.2, t_12.6_ = −4.076, *p* = 0.001) and 2-caves treatment (LMM, status: −17.1±4.2, t_12.6_ = −4.076, *p* = 0.001), while no significant difference was found in the 3-caves treatment (LMM, status: −5.13±7.43, t_8.7_ = −0.690, *p* = 0.508, **Figure 3b**). Cave number treatment did not affect testosterone levels (*p* = 0.19, **Table 1**) after controlling for the effect of social status (*p* < 0.001, **Table 1**). Within subordinate males, there was an increase in testosterone levels with cave numbers, but this effect was not significant (LM, caves: 5.318±2.776, t = 1.915, *p* = 0.067). Cave number treatment did not affect testosterone levels within dominant males (LM, caves: 0.053±2.824, t = 0.019, *p* = 0.985).

#### Behavior

We calculated the average rate of different behaviors in each compartment across the five weekly five-minute observations. Dominant males engaged more frequently in chases, aggressive display behaviors, courtship, cave visits, and foraging than subordinate males (all *p values* < 0.001, **Table 2**). In contrast, subordinate males displayed significantly more fleeing behavior than dominant males (*p* < 0.001) (**Figure 4b**). None of these social status effects were influenced by cave number treatment with the exception of chase rate, which varied across cave number treatments as indicated by a significant interaction effect between social status and cave number treatment (*p* = 0.044, **Figure 4a**). The interaction was driven by a slight reduction in the difference in chasing behavior between dominant and subordinate individuals (**Figure 4a**).

**Figure 4:**
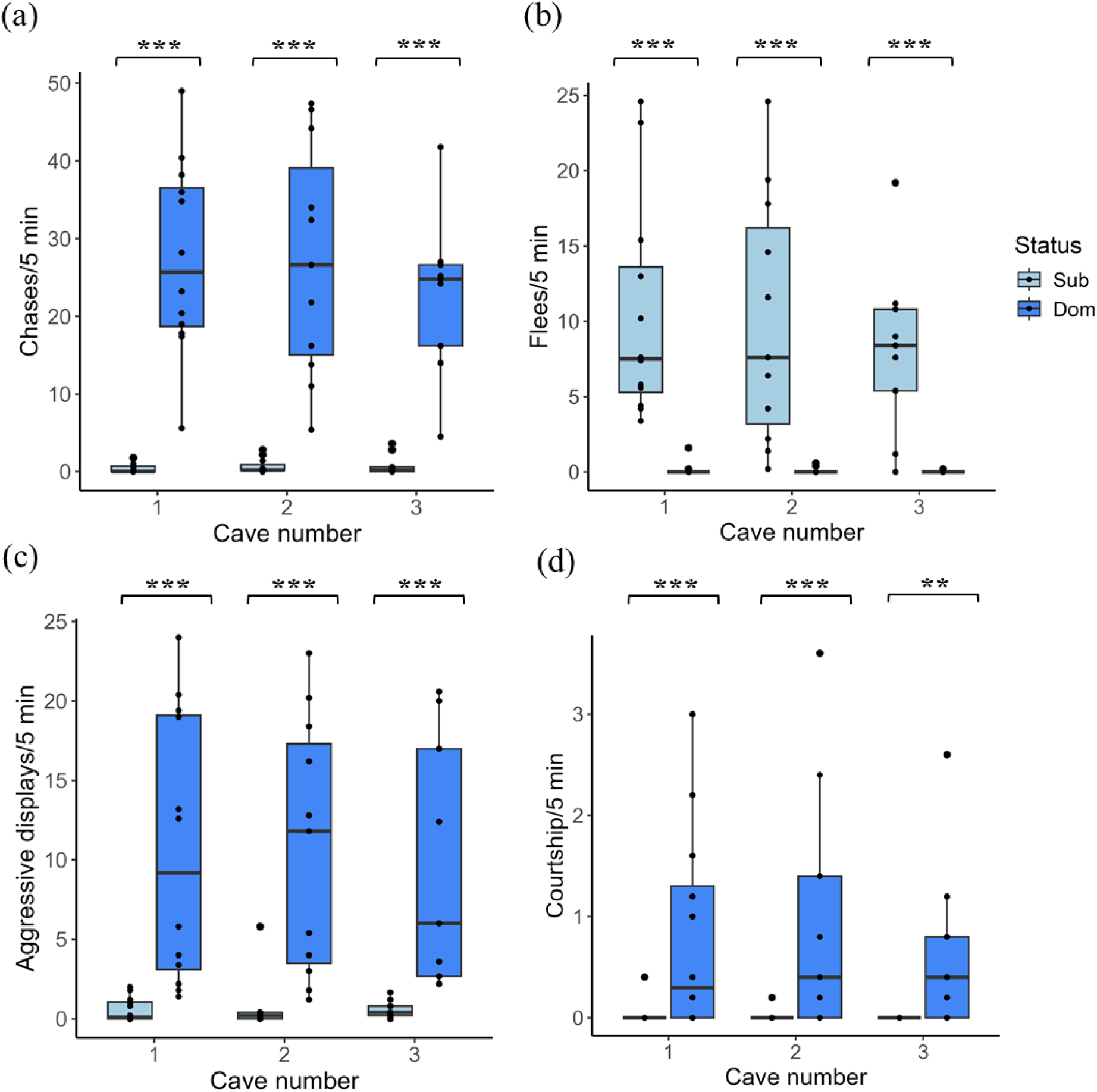
(a) Chases, (b) flees, (c) aggressive displays, (d) courtship behaviors of subordinate (Sub) and dominant (Dom) males for 1-cave, 2-caves and 3-caves treatments. Data was averaged across 5-week period. Boxes represent 25th to 75th percentiles. Error bars represent data range. \*\**p* < 0.01, \*\*\**p* < 0.001.

**Table 2:** Relationship between behaviors and cave treatments in dominant and subordinate *Astatotilapia burtoni* males. We analyzed the data using generalized linear mixed models (with negative binomial distribution) with social status as a fixed factor and group identity as a random factor. Shown are the interaction effects and main effects. Significant effects (*p* < 0.05) are indicated in bold.

| Behavior | Effect | Estimate | z value | p value |
| --- | --- | --- | --- | --- |
| Chases | Cave | -0.036±0.113 | -0.315 | 0.753 |
|  | Status | 3.649±0.178 | 20.415 | **<0.001\*\*\*** |
|  | Cave*status | -0.434±0.216 | -2.011 | **0.044 \*** |
| Aggressive displays | Cave | -0.0008±0.184 | -0.004 | 0.996 |
|  | Status | 2.875±0.178 | 16.15 | **<0.001\*\*\*** |
|  | Cave*Status | -0.089±0.210 | -0.425 | 0.671 |
| Courtship | Cave | -0.052±0.304 | -0.171 | 0.864 |
|  | Status | 3.639±0.721 | 5.05 | **<0.001\*\*\*** |
|  | Cave*Status | 0.615±1.069 | 0.575 | 0.565 |
| Cave visits | Cave | 0.211±0.110 | 1.915 | 0.055 |
|  | Status | 3.745±0.456 | 8.219 | **<0.001\*\*\*** |
|  | Cave*Status | -0.677±0.617 | -1.099 | 0.272 |
| Flees | Cave | -0.070±0.155 | -0.451 | 0.652 |
|  | Status | -4.414±0.418 | -10.554 | **<0.001\*\*\*** |
|  | Cave*Status | -0.309±0.549 | -0.562 | 0.574 |
| Foraging | Cave | -0.117±0.115 | -1.022 | 0.306 |
|  | Status | 2.214±0.214 | 10.336 | **<0.001\*\*\*** |
|  | Cave*Status | -0.243±0.262 | -0.928 | 0.353 |

## Discussion

We studied how environmental enrichment influences animal welfare and social dominance relationships in the cichlid *A. burtoni*. We found that subordinate males exhibited more injuries, lower specific growth rates and reduced body condition compared to dominant males, but none of these markers of welfare were significantly influenced by the cave number treatment within subordinate males. Consistent with previous findings, dominant males exhibited higher rates of territorial behavior, higher relative gonad mass (gonadosomatic index (GSI)), and higher levels of testosterone than subordinate males. However, differences in testosterone levels, GSI and chases were diminished between dominant and subordinate males as environmental complexity increased. These findings suggest that environmental enrichment, when provided as an enrichment structure that could be monopolized by a single dominant male, has little to no effect on the welfare of subordinate individuals but diminishes the physiological and behavioral differentiation between the two social states.

The lack of a positive effect of cave numbers on welfare in subordinates was somewhat surprising. Visual inspection of the data suggested a positive effect of cave number on body condition in subordinate males, but this pattern failed to reach significance. A more enriched environment may provide more hiding space or limit visibility (Höjesjö et al., 2004; Baird et al., 2006; Kadry and Barreto, 2010), which could benefit subordinates by avoiding direct encounters with dominant fish (Barley and Coleman, 2010). However, in our study, subordinates were aggressively excluded from the cave(s) by the dominant male across all three cave treatments. Even though cave number attenuated the differences in chase rate between dominant and subordinate males, the latter were still subjected to significant rates of chases, as indicated by high rates of fleeing in all three treatments. This sustained high level of aggression regardless of enrichment treatment could potentially explain the lack of a clear effect of cave numbers on our markers of welfare.

As expected, dominant males were more aggressive, had higher testosterone levels and greater relative gonad mass values than subordinate males across all cave treatments. However, the differences in testosterone levels between dominant and subordinate males was not significant in the 3-caves treatment, possibly due to the marginal, though non-significant, increase in testosterone among subordinate males with increasing cave number. Regarding chasing behavior, a significant interaction effect indicated that differences between dominant and subordinate males diminished as cave number increased. Subordinate males are known to exhibit elevated testosterone levels during male-male interactions (Oliveira et al., 2002) and often engage in increased agonistic behavior during periods of social instability to attempt social ascension (Desjardins et al., 2012; Maguire et al., 2021). Elevated testosterone in subordinate males may serve as an anticipatory mechanism in preparation of social ascent and future reproductive opportunities (Huffman et al., 2012; Maruska and Fernald, 2013). The increase in cave number, despite being clumped and defendable by only one dominant male, may have still enhanced the perceived opportunity for social ascent, thereby reducing stress-related suppression of reproductive hormones. The reduced physiological and behavioral differences between dominant and subordinate males with increasing cave number could hinder studies where distinct differences in behavior and physiology between social states are required (Alward et al., 2019; Friesen et al., 2022; Maruska et al., 2022; Dijkstra et al., 2024) or where subordinate males need to exhibit social suppression of the reproductive axis, which is typically associated with low androgen levels (Maruska and Fernald, 2013).

Environmental enrichment is widely used to enhance the welfare of laboratory animals, with numerous studies demonstrating its benefits across various experimental settings (Simpson and Kelly, 2011; Zhang et al., 2022). However, laboratories investigating status or rank-based behaviors may encounter variability in experimental outcomes due to the influence of enrichment on social dynamics. While our findings provide valuable insights, some limitations must be considered. In our experimental design, structural complexity was positively linked to structure size. The size of defendable structures may enhance resource holding potential and are preferentially occupied in some cichlid species (Dijkstra et al., 2008). In addition, we used halved terracotta pots as defendable structures; however, using different structures [such as PVC (Dijkstra et al., 2008; Näslund and Johnsson, 2016)] or altering the placement or orientation of defendable structures may yield varying results. Future research should examine how specific structural properties (complexity, size, and placement) shape aggression and welfare in cichlids as well as other social fish species.

Our findings suggest that enhancing enrichment by increasing the number of caves in a clumped manner has little to no positive effect on animal welfare in subordinate cichlid males. This is an important contribution as we were able to isolate the effect of altering the complexity of a clumped structure that could only be defended by one dominant cichlid male. Furthermore, we found that increasing environmental complexity reduced behavioral and physiological differences between dominant and subordinate individuals. Whether these status-specific profiles are desirable or not depends on the goal of the study, but it needs to be taken into account when developing enrichment strategies for laboratory animal species.

## Acknowledgements

We thank the members of the Dijkstra Lab for their assistance with the experiment and data collection. This research was supported by grants from the American Association for Laboratory Animal Science (Grants for Laboratory Animal Science) to PDD and JK, the National Institute of General Medical Sciences (NIGMS, R15GM150286 to PDD), and the National Science Foundation (grant number 2444902 to PDD).

## Notes

### Competing Interest Statement

The authors have declared no competing interest.

## References

Alcazar, R. M., Becker, L., Hilliard, A. T., Kent, K. R. and Fernald, R. D. (2016). Two types of dominant male cichlid fish: Behavioral and hormonal characteristics. Biol Open 5, 1061–1071.

Alward, B. A., Hilliard, A. T., York, R. A. and Fernald, R. D. (2019). Hormonal regulation of social ascent and temporal patterns of behavior in an African cichlid. Horm Behav 107, 83–95.

Alward, B. A., Laud, V. A., Skalnik, C. J., York, R. A., Juntti, S. A. and Fernald, R. D. (2020). Modular genetic control of social status in a cichlid fish. Proc Natl Acad Sci U S A 117, 28167–28174.

Bagenal, T. B. and Ricker, W. E. (1978). Methods for assessment of fish production in fresh waters −3d ed. Blackwell Scientific; Distributed by J.B. Lippincott.

Baird, H. P., Patullo, B. W. and Macmillan, D. L. (2006). Reducing aggression between freshwater crayfish (*Cherax destructor* Clark: Decapoda, Parastacidae) by increasing habitat complexity. Aquac Res 37, 1419–1428.

Barley, A. J. and Coleman, R. M. (2010). Habitat structure directly affects aggression in convict cichlids *Archocentrus nigrofasciatus*. Curr Zool 56, 52–56.

Basquill, S. P. and Grant, J. W. (1998). An increase in habitat complexity reduces aggression and monopolization of food by zebra fish (*Danio rerio*). Can J Zool 76, 770–772.

Bates, D., Mächler, M., Bolker, B. and Walker, S. (2015). Fitting Linear Mixed-Effects Models Using lme4. J Stat Softw 67, 1–48.

Benefiel, A. C., Dong, W. K. and Greenough, W. T. (2005). Mandatory “enriched” housing of laboratory animals: the need for evidence-based evaluation. ILAR J 46, 95–105.

Blanchet, S., Dodson, J. J. and Brosse, S. (2006). Influence of habitat structure and fish density on Atlantic salmon *Salmo salar* L. territorial behaviour. J Fish Biol 68, 951–957.

Bolger, T. and Connolly, P. L. (1989). The selection of suitable indices for the measurement and analysis of fish condition. J Fish Biol 34, 171–182.

Border, S. E., Deoliveira, G. M., Janeski, H. M., Piefke, T. J., Brown, T. J. and Dijkstra, P. D. (2019). Social rank, color morph, and social network metrics predict oxidative stress in a cichlid fish. Behav Ecol 30, 490–499.

Brooks, M. E., Kristensen, K., Van Benthem, K. J., Magnusson, A., Berg, C. W., Nielsen, A., Skaug, H. J., Mächler, M. and Bolker, B. M. (2017). glmmTMB Balances Speed and Flexibility Among Packages for Zero-inflated Generalized Linear Mixed Modeling. R J 9, 378.

Culbert, B. M., Border, S. E., Fialkowski, R. J., Bolitho, I. and Dijkstra, P. D. (2023). Social status influences relationships between hormones and oxidative stress in a cichlid fish. Horm Behav 152, 105365.

Desjardins, J. K., Hofmann, H. A. and Fernald, R. D. (2012). Social Context Influences Aggressive and Courtship Behavior in a Cichlid Fish. PLoS One 7, e32781.

Dijkstra, P. D., van der Zee, E. M. and Groothuis, T. G. G. (2008). Territory quality affects female preference in a Lake Victoria cichlid fish. Behav Ecol Sociobiol 62, 747–755.

Dijkstra, P. D., Fialkowski, R. J., Bush, B., Wong, R. Y., Moore, T. I. and Harvey, A. R. (2024). Oxidative stress in the brain is regulated by social status in a highly social cichlid fish. Front Behav Neurosci 18, 1477984.

du Sert, N. P., Hurst, V., Ahluwalia, A., Alam, S., Avey, M. T., Baker, M., Browne, W. J., Clark, A., Cuthill, I. C., Dirnagl, U., et al. (2020). The arrive guidelines 2.0: Updated guidelines for reporting animal research. PLoS Biol 18, 1–12.

Eason, P. K. and Stamps, J. A. (1992). The effect of visibility on territory size and shape. Behav Ecol 3, 166–172.

Fernald, R. D. and Hirata, N. R. (1979). The Ontogeny of Social Behavior and Body Coloration in the African Cichlid Fish *Haplochromis burtoni*. Z Tierpsychol 50, 180–187.

Fialkowski, R., Aufdemberge, P., Wright, V. and Dijkstra, P. (2021). Radical change: temporal patterns of oxidative stress during social ascent in a dominance hierarchy. Behav Ecol Sociobiol 75, 43.

Fialkowski, R. J., Border, S. E., Bolitho, I. and Dijkstra, P. D. (2022). Social dominance and reproduction result in increased integration of oxidative state in males of an African cichlid fish. Comp Biochem Physiol A Mol Integr Physiol 269, 111216.

Friesen, C. N., Maclaine, K. D. and Hofmann, H. A. (2022). Social status mediates behavioral, endocrine, and neural responses to an intruder challenge in a social cichlid, *Astatotilapia burtoni*. Horm Behav 145, 105241.

Galhardo, L., Correia, J. and Oliveira, R. (2008). The effect of substrate availability on behavioural and physiological indicators of welfare in the African cichlid (*Oreochromis mossambicus*). Anim Welf 17, 239–254.

Giles, J. M., Whitaker, J. W., Moy, S. S. and Fletcher, C. A. (2018). Effect of environmental enrichment on aggression in BALB/cJ and BALB/cByJ mice monitored by using an automated system. J Am Assoc Lab Anim Sci 57, 236–243.

Höjesjö, J., Johnsson, J. and Bohlin, T. (2004). Habitat complexity reduces the growth of aggressive and dominant brown trout (*Salmo trutta*) relative to subordinates. Behav Ecol Sociobiol 56, 286–289.

Hoyle, I., Oidtmann, B., Ellis, T., Turnbull, J., North, B., Nikolaidis, J. and Knowles, T. G. (2007). A validated macroscopic key to assess fin damage in farmed rainbow trout (*Oncorhynchus mykiss*). Aquaculture 270, 142–148.

Huffman, L. S., Mitchell, M. M., O’Connell, L. A. and Hofmann, H. A. (2012). Rising StARs: Behavioral, hormonal, and molecular responses to social challenge and opportunity. Horm Behav 61, 631–641.

Ibi, M., Liu, J., Arakawa, N., Kitaoka, S., Kawaji, A., Matsuda, K. ichi, Iwata, K., Matsumoto, M., Katsuyama, M., Zhu, K., et al. (2017). Depressive-like behaviors are regulated by NOX1/NADPH oxidase by redox modification of NMDA receptor 1. J Neurosci 37, 4200–4212.

Imre, I., Grant, J. W. and Keeley, E. R. (2002). The effect of visual isolation on territory size and population density of juvenile rainbow trout (*Oncorhynchus mykiss*). Can J Fish Aquat Sci 59, 303–309.

Jobling, M. (1983). Growth studies with fish—overcoming the problems of size variation. J Fish Biol 22, 153–157.

Kadry, V. O. and Barreto, R. E. (2010). Environmental enrichment reduces aggression of pearl cichlid, Geophagus brasiliensis, during resident-intruder interactions. Neotrop Ichthyol 8, 329–332.

Kaliste, E. K., Mering, S. M. and Huuskonen, H. K. (2006). Environmental modification and agonistic behavior in NIH/S male mice: Nesting material enhances fighting but shelters prevent it. Comp Med 56, 202–208.

Lehmann, M. L., Weigel, T. K., Poffenberger, C. N. and Herkenham, M. (2019). The behavioral sequelae of social defeat require microglia and are driven by oxidative stress in mice. J Neurosci 39, 5594–5605.

Lidster, K., Owen, K., Browne, W. J. and Prescott, M. J. (2019). Cage aggression in group-housed laboratory male mice: an international data crowdsourcing project. Sci Rep 9, 1–12.

Maguire, S. M., DeAngelis, R., Dijkstra, P. D., Jordan, A. and Hofmann, H. A. (2021). Social network dynamics predict hormone levels and behavior in a highly social cichlid fish. Horm Behav 132, 104994.

Mandrekar, K. and Oldfield, R. G. (2008). Prior residency and social experience in contests between similar-sized juvenile black Midas cichlids, Amphilophus astorquii. Aqua 14, 141–148.

Maruska, K. P. and Fernald, R. D. (2011). Social regulation of gene Expression in the hypothalamic-pituitary-gonadal axis. Physiol J 26, 412–423.

Maruska, K. P. and Fernald, R. D. (2013). Social Regulation of Male Reproductive Plasticity in an African Cichlid Fish. Integr Comp Biol 53, 938–950.

Maruska, K. P. and Fernald, R. D. (2018). *Astatotilapia burtoni*: a model system for analyzing the neurobiology of Behavior. ACS Chem Neurosci 9, 1951–1962.

Maruska, K. P., Anselmo, C. M., King, T., Mobley, R. B., Ray, E. J. and Wayne, R. (2022). Endocrine and neuroendocrine regulation of social status in cichlid fishes. Horm Behav 139, 105110.

Näslund, J. and Johnsson, J. I. (2016). Environmental enrichment for fish in captive environments: Effects of physical structures and substrates. Fish Fish 17, 1–30.

Noble, C., Kadri, S., Mitchell, D. F. and Huntingford, F. A. (2008). Growth, production and fin damage in cage-held 0+ Atlantic salmon pre-smolts (*Salmo salar* L.) fed either a) on-demand, or b) to a fixed satiation–restriction regime: Data from a commercial farm. Aquaculture 275, 163–168.

O’Connell, L. A. and Hofmann, H. A. (2012). Social status predicts how sex steroid receptors regulate complex behavior across levels of biological organization. Endocrinology 153, 1341–51.

Oldfield, R. G. (2011). Aggression and Welfare in a Common Aquarium Fish, the Midas Cichlid. J Appl Anim Welf Sci 14, 340–360.

Oliveira, R. F., Hirschenhauser, K., Carneiro, L. A. and Canario, A. V. M. (2002). Social modulation of androgen levels in male teleost fish. Comp Biochem Physiol B Biochem Mol Biol 132, 203–215.

Patki, G., Solanki, N., Atrooz, F., Allam, F. and Salim, S. (2013). Depression, anxiety-like behavior and memory impairment are associated with increased oxidative stress and inflammation in a rat model of social stress. Brain Res 1539, 73–86.

Piefke, T. J., Bonnell, T. R., DeOliveira, G. M., Border, S. E. and Dijkstra, P. D. (2021). Social network stability is impacted by removing a dominant male in replicate dominance hierarchies of a cichlid fish. Anim Behav 175, 7–20.

Ruberto, T., Swaney, W. T. and Reddon, A. R. (2024). Submissive behavior is affected by territory structure in a social fish. Curr Zool 70, 803–809.

Simpson, J. and Kelly, J. P. (2011). The impact of environmental enrichment in laboratory rats—Behavioural and neurochemical aspects. Behav Brain Res 222, 246–264.

Smith, C. (2011). Good fences make good neighbours: the role of landmarks in territory partitioning in the rose bitterling (*Rhodeus ocellatus*). Behaviour 148, 233–246.

Snyder-Mackler, N., Burger, J. R., Gaydosh, L., Belsky, D. W., Noppert, G. A., Campos, F. A., Bartolomucci, A., Yang, Y. C., Aiello, A. E., O’Rand, A., et al. (2020). Social determinants of health and survival in humans and other animals. Science (1979) 368, eaax9553.

Sundbaum, K. and Näslund, I. (1998). Effects of woody debris on the growth and behaviour of brown trout in experimental stream channels. Can J Zool 76, 56–61.

Tallent, B. R., Law, L. M. and Lifshitz, J. (2024). Partially divided caging reduces overall aggression and anxiety which may indicate improved welfare in group housed male C57BL/6J mice. BMC Vet Res 20, 69.

Theil, J. H., Ahloy-Dallaire, J., Weber, E. M., Gaskill, B. N., Pritchett-Corning, K. R., Felt, S. A. and Garner, J. P. (2020). The epidemiology of fighting in group-housed laboratory mice. Sci Rep 10, 1–10.

Wilkes, L., Owen, S. F., Readman, G. D., Sloman, K. A. and Wilson, R. W. (2012). Does structural enrichment for toxicology studies improve zebrafish welfare? Appl Anim Behav Sci 139, 143–150.

Young, R. J. (2013). Environmental Enrichment for Captive Animals. Wiley.

Zhang, Z., Gao, L. and Zhang, X. (2022). Environmental enrichment increases aquatic animal welfare: A systematic review and meta-analysis. Rev Aquac 14, 1120–1135.

