## Supplementary File for "The effect of environmental enrichment on social dominance and welfare in a cichlid fish"

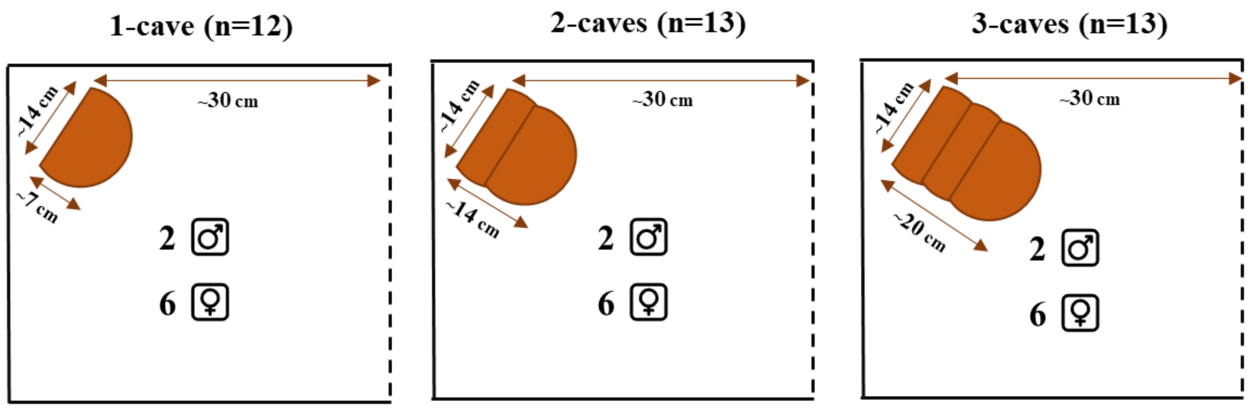


**Supplementary figure 1:** Top view of cave placement in experimental tanks. Treatments included 1-cave (n = 12), 2-caves (n = 13), and 3-caves (n = 13) setups. Caves were positioned away from the central divider and approximately 2 cm from the tank wall. Each compartment housed 2 males and 6 females. A neighboring compartment with an identical group composition was visible through a perforated screen (dashed line).

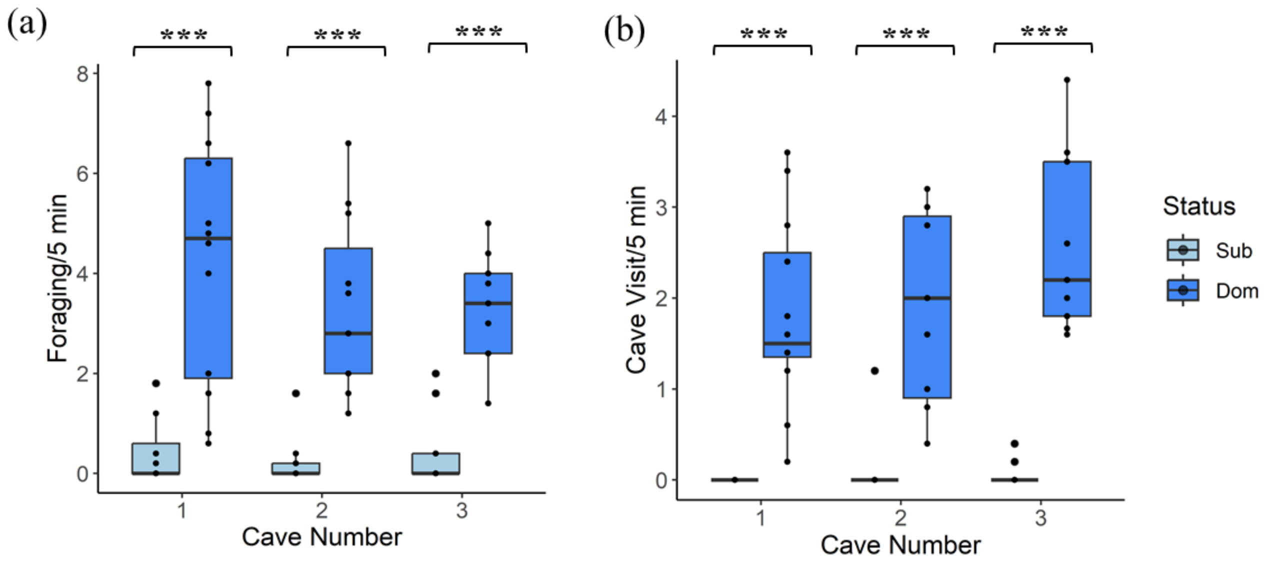


**Supplementary figure 2:** (a) Foraging, (b) cave visits behaviors of subordinate (Sub) and dominant (Dom) males for 1-cave, 2-caves and 3-caves treatments. Data was averaged across 5-week period. Boxes represent 25th to 75th percentiles. Error bars represent data range. ***$p < 0.001$.
